# Soluble CD93 is an Apoptotic Cell Opsonin Recognized by the α_x_β_2_ Integrin

**DOI:** 10.1101/341933

**Authors:** Jack W.D. Blackburn, Darius H.C. Lau, Jessica Ellins, Angela Kipp, Emily N. Pawlak, Jimmy D. Dikeakos, Bryan Heit

## Abstract

Efferocytosis – the phagocytic removal of apoptotic cells – is essential for the maintenance of homeostasis and prevention of the inflammatory and autoimmune diseases which can follow the lysis of uncleared apoptotic cells. CD93 is a transmembrane glycoprotein previously implicated in efferocytosis and angiogenesis, and upon mutation, results in the onset of efferocytosis-associated diseases such as atherosclerosis and rheumatoid arthritis. CD93 is produced as a cell surface protein which is shed as soluble CD93, but it is unknown how CD93 mediates efferocytosis or whether its efferocytic activity is mediated by the soluble or membrane-bound form. Herein, we demonstrate that the membrane bound form of CD93 has no phagocytic, efferocytic, or tethering activity, whereas soluble CD93 potently opsonizes apoptotic cells but not a broad range of Gram-Negative, Gram-Positive or fungal microorganisms. Using mass spectrometry, we identified the α_x_β_2_ integrin as the receptor required for soluble CD93-mediated efferocytosis, and via deletion mutagenesis determined that soluble CD93 binds to apoptotic cells via its C-Type Lectin-Like domain, and to α_x_β_2_ by its EGF-like repeats. This bridging of apoptotic cells to the α_x_β_2_ integrin markedly enhanced efferocytosis by macrophages, and could be abrogated by knockdown of α_x_β_2_ integrin. Combined, these data elucidate the mechanism by which CD93 regulates efferocytosis and identify a previously unreported opsonin-receptor system utilized by the immune system for the efferocytic clearance of apoptotic cells.

## Introduction

CD93 is a 120kDa, type-1 transmembrane glycoprotein expressed on the surface of endothelial cells, stem cells, platelets, circulating myeloid cells, and on maturing B cells (1–6). Part of the group XIV C-type lectin family, CD93 shares a common structure with the other group XIV family proteins thrombomodulin, CLEC14A and CD248: an N-terminal C-type lectin-like domain (CTLD), five EGF-like repeats, a heavily glycosylated mucin domain, a transmembrane helix, and a small and highly cationic cytosolic domain (7). A soluble form of CD93 (sCD93) comprised of the CTLD, EGF-like domain and a portion of the mucin domain, can be detected in human plasma (4). The function of sCD93 remains poorly elucidated, however sCD93 serum levels are increased during inflammatory conditions such as atherosclerosis and in autoimmune disorders such as rheumatoid arthritis (8, 9). CD93 was originally identified as a receptor for C1q (7, 10), based on observations that CD93 enhanced phagocytosis in a C1q-dependent manner (10–14). These results are controversial, as subsequent studies demonstrated that CD93 neither interacts with C1q, nor contributes to the C1q dependent enhancement of phagocytosis; findings later confirmed in knockout mice (15, 16). Furthermore, single nucleotide polymorphisms in CD93 are strongly associated with diseases characterized by impaired apoptotic cell clearance, such as atherosclerosis and systemic lupus erythematosus (17–19), suggesting that CD93 may contribute to efferocytosis – the phagocytic removal of apoptotic cells. Indeed, CD93 was first identified by thiohydantoin C-terminal sequencing of a protein immunoprecipitated by an antibody known to block the efferocytic activity of monocyte-like U937 cells (11). Moreover, CD93^−/−^ mice have significant defects in apoptotic cell clearance compared to strain matched controls, and CD93 polymorphisms predispose mice and humans to autoimmune and inflammatory disorders characterized by failed efferocytosis (7, 16–18). As one example, patients homozygous for a P^541^S missense mutation in the CD93 mucin domain have a 26% increase in their risk of coronary heart disease; a disease resulting from the accumulation of uncleared apoptotic cells beneath the cardiac vasculature (17, 20).

Although the above studies indicate that CD93 is involved in efferocytosis, it remains unclear how CD93 mediates this process. While some studies have suggested that CD93 functions as a membrane-bound receptor which directly induces the engulfment of apoptotic cells (18, 19, 21), CD93 lacks the ITAM signaling motifs normally associated with phagocytic/efferocytic receptors. Instead, CD93 has a short intracellular domain enriched in the cationic amino acids lysine and arginine, and associates with moesin and GIPC – signaling molecules not normally associated with phagocytosis (22, 23). Although CD93 does not interact with classical mediators of phagocytosis, moesin-CD93 interactions are regulated by phosphatidylinositol-3-kinase signaling similar to that present during phagocytosis. Moreover, the intracellular domain of CD93 can be phosphorylated via a dystroglycan-dependent pathway that results in rearrangement of the cell cytoskeleton (23, 24). Similarly, GIPC is involved in receptor-mediated endocytosis, and therefore may be co-opted for the uptake of larger cargos (25). While not conclusively demonstrated, it is possible that these pathways may represent a novel signaling pathway that can drive the cytoskeletal rearrangements required for the engulfment of apoptotic cells. A second possibility is that CD93 acts as a tethering receptor – e.g. CD93 stabilizes interactions between phagocytes and apoptotic cells, thereby prolonging contact with other efferocytic receptors. Indeed, CD93 is recognized to play a role in intercellular adhesion (3, 15, 24, 26, 27), consistent with a tethering function. In conflict with these two models is the observation that sCD93-rich peritoneal lavage fluid from wild-type, but not CD93^−/−^ mice, greatly enhanced efferocytosis by bone-marrow derived CD93^−/−^ macrophages (28). This data indicates that the efferocytic capacity of CD93 is localized to the soluble form, thereby supporting a third hypothesis – that CD93 does not act directly as an efferocytic receptor, and instead acts as a soluble opsonin.

Clearly, there is data supporting three distinct and mutually exclusive models of CD93 efferocytic function. In this study, we directly asses these three models of CD93 function, using in vitro models of binding, efferocytosis and opsonization. These experiments disfavor the efferocytic receptor and tethering receptor models of CD93 function, and instead demonstrate a strong opsonic effect of sCD93 against apoptotic cells but not against a panel of microorganisms. Using mass-spectrometry we then identified α_x_β_2_ integrin as the efferocytic receptor which recognizes sCD93 opsonized targets, with apoptotic cells “bridged” by the CD93 CTLD domain to α_x_β_2_ integrin on phagocytes via the CD93 EGF-like domain. This represents a previously undescribed opsonin-receptor system which aids in the efferocytosis of apoptotic cell by phagocytes such as macrophages.

## Materials and Methods

### Materials

DH5a *Escherichia coli* was a gift from Dr. John McCormick (University of Western Ontario), *Rhodotorula minuta* and *Candida glabrate* were gifts from Dr. Andre Lachance (University of Western Ontario). *Escherichia coli* K29^+^ and K29^−^, and *Burkholderia cenocepacia* were gifts from Dr. Susan Koval (University of Western Ontario), *Micrococcus luteus* T-18 and *Staphylococcus pyrogenes* 71-679 were gifts from Dr. Jeremy Burton (University of Western Ontario), *Staphylococcus aureus* UWO256 and *Pseudomonas aeruginosa* UWO533 were from the University of Western Ontario microbiology teaching bank. The 12CA5 hybridoma (mouse-anti-HA) and THP-1 monocytes were gifts from Dr. Joe Mymryk (University of Western Ontario). FcγRI and TIM4 expression constructs were a gift from Dr. Sergio Grinstein (Hospital for Sick Children, Toronto). Saccharomyces cerevesea WLP001 was purchased from White Labs (San Diego, CA), Jurkat, J774.2 and CHO cells, and all fluorescent secondary antibodies were purchased from Cedarlane (Mississauga, Canada). Goat-anti-Human CD93 polyclonal serum was purchased from R&D Systems (Oakville, Canada). Roswell Park Memorial (RPMI), Dulbecco’s Modified Eagle Medium (DMEM), serum-free CHO-100 medium, serum-free hybridoma medium, Trypsin-EDTA, and Fetal Bovine Serum (FBS) were purchased from Wisent (Saint-Jean-Baptiste, Canada). #1.5 thickness round cover slips, glass slides and 16% paraformaldehyde (PFA) were purchased from Electron Microscopy Sciences (Hatfield, Pennsylvania). DTT, human IgG and cysteine was purchased from Sigma-Aldrich (Oakville, Canada). GenJet Plus, Polyjet and all DNA purification kits were purchased from Frogga Bio (North York, Canada). DAPI, permafluor mounting medium, 30 kDa cut-off spin concentrators, FITC anti-human CD3, and Phusion DNA polymerase, and N-Ethyl-N’-(3-dimethylaminopropyl)carbodiimide were purchased from Thermo Scientific (Mississauga, Canada). Restriction enzymes and T4 DNA ligase was purchased from New England Biolabs (Whitby, Canada). Magnetic 1 µm diameter carboxy-reactive beads, 3 µm diameter silica beads, 3 µm diameter P(S/DVB) polystyrene beads and 1 μm diameter Promag carboxy magnetic microspheres were purchased from Bangs Laboratories (Fishers, Indiana). Phosphatidylcholine, phosphatidylserine and biotin-phosphatidylethanolamine were purchased from Avanti Polar Lipids, Inc. (Alabaster, Alabama). Anti-β_1_ (clone AIIB2), anti-β_1_ (clone H52) and anti-CD44 (clone H4C4) were purchased from the Developmental Studies Hybridoma Bank (University of Iowa). Anti-α_x_ (clone BU15) and all immunoblotting supplies were purchased from Bio-Rad Canada (Montreal, Canada). All other chemicals and bacterial culture media were purchased from Canada BioShop (Mississauga, Canada). Matlab software was purchased from MathWorks (Natick, Massachusetts). Prism software was purchased from Graphpad (La Jolla, California). FIJI is just ImageJ (FIJI) was downloaded from https://fiji.sc/ (29).

### Cell Culture

CHO and HEK 293T cells were cultured in Ham’s F12 medium supplemented with 10% FBS, and split upon reaching confluency by washing once with phosphate-buffered saline (PBS, 137 mM NaCl, 10 mM Na_2_HPO_4_) and detached by the addition of Trypsin-EDTA. J774.2 cells were grown in DMEM supplemented with 10% FBS, and split by detaching cells with a cell scraper. Detached CHO, HEK 293T, and RAW264.7 cells were then passaged by diluting 1:10 into fresh media. THP-1, Jurkat and 12CA5 cells were grown in RPMI supplemented with 10% FBS and DMEM supplemented with 10% hybridoma grade FBS respectively, and split by diluting 1:10 upon reaching a density of ∼2 × 10^6^ cells/ml. For phagocytosis/efferocytosis experiments, ∼1 × 10^5^ J774.2 or CHO cells were split onto #1.5 thickness, 18 mm diameter circular coverslips placed into the wells of a 12-well plate. If required, 18-24 hours later the cells were transfected with 1 μg DNA/well using FugeneHD (J774.2) or GenJet (CHO) as per manufacturers’ instruction. To generate THP-1 derived macrophages, 1 × 10^5^ THP-1 cells were placed into the well of a 12-well plate, and the media supplemented with 10 ng/mL PMA, and the resulting macrophages used 72 hours later. To generate apoptotic cells, 5 ml of Jurkat cell culture at ∼2 × 10^6^ cells/mL was resuspended in 1.5 mL of serum-free RPMI supplemented with 10 μM staurosporine following a 350×g/5 min centrifugation, and incubated overnight at 37°C. Apoptotic cells were suspended in PBS using a 1,500×g/1 min centrifugation, labeled for 20 min with a 1:500 dilution of Cell Proliferation Dye eFluor 670, washed 2× with PBS, and suspended in serum-free RPMI for immediate use.

### Primary Human Macrophage Culture

PBMCs were suspended in RPMI-1640 supplemented with 10% FBS and 1% antibiotic–antimycotic solution at ∼2 × 10^6^ cells/mL. Monocytes were separated by adhesion to glass by placing 200 μl or 1.0 ml of the suspension on sterile 18 mm coverslips or on 35 mm diameter gelatin-coated 6-well tissue culture plates. Non-adherent cells were removed after a 1 h/37 °C incubation with two PBS washes. Monocytes were differentiated into M0, M1 or M2 polarized macrophages as per our published protocols (30, 31). Briefly, adherent monocytes were cultured for 5 days in 10 ng/ml M-CSF (M0 and M2) or 20 ng/ml GMCSF (M1), followed by 2 days culture in DMEM supplemented with 10% FBS with 10ng/ml M-CSF (M0), 20 ng/ml GM-CSF supplemented with 250 ng/ml *Salmonella* lipopolysaccharide and 10 ng/ml INFγ (M1), or 10 ng/ml M-CSF and 10 ng/ml IL-4 (M2).

### Antibody and Fab Preparation

12CA5 cells were grown to ∼2 × 10^6^ cells/mL, collected with a 350×g/10 min centrifugation, suspended in 60 ml of serum-free hybridoma media, and cultured for an additional 4 days. Cells and particulates were removed by a 3,000×g/30 min centrifugation, and the antibodies concentrated and media exchanged to TBS (10mM tris + 150mM NaCl), using a 10 kDa cutoff spin concentrator. For full-length antibody, the resulting concentrate was diluted to 1 mg/ml in TBS with 20% glycerol and stored at −20°C. Fab’s were prepared by diluting the concentrate to 5 mg/ml in TBS with 20mM cysteine-HCl at pH = 7.0; 0.5 ml of immobilized papain was added per 1 ml of antibody, incubated overnight on a rotator at 37°C, the reaction terminated by the addition of an equal volume of TBS pH = 7.5, and the papain removed by a 21,000×g/5 min centrifugation. Fc fragments were depleted by the addition of 0.5 ml of immobilized protein A, followed by a 4 hr room-temperature incubation, and separation of the Fc fragments by a 21,000×g/5 min centrifugation. Fabs were then purified to high purity (>90%) by FPLC using an Enrich SEC650 size-exclusion column and a BioRad NGC FPLC platform. Purified Fab fragments were suspended at 0.5 mg/ml in TBS with 20% glycerol and stored at −20°C.

### Preparation of CD93 Expression and shRNA Vectors and Recombinant sCD93

The human CD93 ORF was ordered from the Harvard PlasmID Database. Expression vectors were generated using either restriction digest or Gibson assembly using the primers and enzymes indicated in Supplemental Table 1. All PCR reactions were performed using Phusion DNA polymerase with 35-40 cycles of 96°C/30 sec melting, 65-63°C/30 sec annealing and 72°C/90 sec elongation. Restriction digests and ligations were performed for an hour at 37°C and 20°C, respectively. Gibson assembly was performed as per manufacturer’s instructions. shRNAs were designed using the Hanon Lab shRNA design webserver (http://cancan.cshl.edu/RNAi_central). shRNA inserts were ordered as complementary single-stranded oligonucleotides which were annealed by mixing the oligos in equimolar amounts in ddH_2_O, heating to 95°C, and then cooling at 2°C/min to 20°C. shRNA inserts and pGFP-C-shLenti vectors were digested with AgeI/BamHI and ligated with T4 DNA ligase. All vector sequences were confirmed with Sanger sequencing. The sCD93 and shRNA vectors were packaged into a VSV-G pseudotyped lentiviral vector by triple transfection of HEK 293T cells with the VSV-G envelope-encoding pMD2.G (2 ng, Addgene #12259), pCMV-DR8.2 (5 ng, Addgene #12263) and 2.5 µg of the lentiviral vector of interest using PolyJet as per the manufacturer’s protocol. Seventy-two hour post transfection, lentivirus was collected in 20% FBS and purified with a 0.2 µm filter, as described previously (32). The resulting virions were stored at −80°C until needed. Virions were used to prepare stably transfected CHO or THP-1 cell lines, using zsGreen or puromycin as a selectable marker. sCD93 proteins were collected by growing the transfected cell lines to confluency in a T75 flask (sCD93) or in 2 wells of a 6-well plate (sCD93-FLAG and deletion mutants), followed by exchanging the medium for 60 ml serum-free CHO100 medium followed by an additional 5 days of culture. The resulting supernatants were collected, cleared with a 3,000×g/30 min centrifugation, sCD93 concentrated and media exchanged to TBS (10mM tris + 150mM NaCl), using a 20 kDa cutoff spin concentrator. sCD93 was then purified to high purity (>95%) by FPLC using an Enrich SEC650 size-exclusion column and a BioRad NGC FPLC platform, suspended at 1 mg/ml in TBS + 20% glycerol and stored at −20°C.

### CD93 Shedding from Human PBMCs

The collection of blood from healthy donors was approved by the Health Science Research Ethics Board of the University of Western Ontario and venipuncture was performed in accordance with the guidelines of the Tri-Council Policy Statement on human research. Blood was collected in heparinized vacuum collection tubes, and 5 ml of blood layered on an equal volume of Lympholyte-poly in a 15 ml centrifuge tube, followed by centrifugation at 300 × g for 30 min at 20 °C. The top band of PBMCs was collected with a sterile transfer pipette, washed once (300 × g, 6 min, 20 °C) with phosphate-buffered saline (PBS, 137 mM NaCl, 2.7 mM KCl, 10 mM Na2HPO4, 1.8 mM KH2PO4), and then suspended in DMEM supplemented with 10% FBS for future experiments. For CD93 shedding experiments, PBMCs or neutrophils were suspended at 1 × 10^6^ cells/mL in serum-free DMEM supplemented with 10 ng/ml GM6001 or 100 ng/ml TAPI-2, and 10 ng/ml PMA, 10 ng/ml LPS or 100 ng/ml TNFα added. Cells were incubated for 60 min at 37°C; for time-courses, 100 μl samples were withdrawn from this culture at desired time-points. Shedding was stopped by cooling the cells on ice and fixating for 20 min with 4% PFA in PBS, and shedding quantified by immunostaining. Shedding following apoptosis was quantified by suspending PBMCs at 1 × 10^6^ cells/mL in serum-free DMEM supplemented with 100 ng/ml TAPI-2 and staurosprorine. A portion of the PBMCs were immediately fixed and immunostained for CD93, and the remaining population incubated for 24 hours at 37°C, followed by fixation and immunostaining for CD93.

### Immmunostaining & Microscopy

Immunolabeling was performed on unfixed cells, or on cells fixed with 4% paraformaldehyde (PFA) in PBS for 20 min at room temperature. For intracellular staining, PFA-fixed cells were permeabilized and blocked with PBS containing 0.1% Triton X-100 and 1% bovine serum albumin (BSA) for 1 hr at room temperature. Primary antibodies were added to the cells in PBS containing 1% BSA at an appropriate dilution (1 μg/ml full-length 12CA5, 0.5 μg/ml 12CA5 Fab or 1:1,000 anti-CD93 polyclonal sera), for 20 min (non-permeabilized) or 1 hr (permeabilized cells) with gentle agitation. Samples were then washed 3 × 15 min in PBS and fluorescently-labeled secondary Fab’s added at a 1:5,000 dilution in PBS containing 1% BSA for 20 min (non-permeabilized) or 1 hr (permeabilized cells) with gentle agitation. If required, cells were counter-stained with 1 μg/ml Hoechst 33342. Samples were then washed 3 × 15 min in PBS and mounted on glass slides with Permafluor mounting media. Samples were imaged on a Leica DMI6000B microscope equipped with 63×/1.40NA and 100×/1.40 NA objectives, photometrics Evolve-512 delta EM-CCD camera, Chroma Sedat Quad filter set and the LAS-X software platform. Z-stacks with 0.19 μm spacing were taken of 30-50 cells/condition, and the resulted images deconvolved using iterative blinded deconvolution in LAS-X, and the resulting images exported for analysis using FIJI (29).

### Bacterial and Fungal Cell Culture & Preparation

*E. coli* was grown in LB medium, *P. aeruginosa, B. cenocepacia, M. luteus, S. pyrogenes,* and *S. aureus* were grown in BHI medium. *R. minuta, C. glabrate,* and *S. cerevisiae* were grown in YPD medium. All microorganisms were grown overnight to stationary phase in 5 ml cultures on a shaking incubator at 37°C (bacteria) or 32°C (fungi). 1 × 10^7^ cells were washed in PBS using 6,000×g 1 min centrifugation, fixed with 4% PFA in PBS for 20 min, and labeled by the addition of a 1:500 dilution of Cell Proliferation Dye eFluor 670 and a 1:500 dilution of NHS-biotin for 30 min at room temperature. After 30 min, the labeling reaction was quenched with LB medium, the organisms washed 3 times with PBS, diluted to 1×10^7^/ml in serum-free DMEM and stored at 4°C until needed.

### Preparation of Apoptotic Cell and Pathogen Mimics

Bead-based mimics were prepared as per our published protocols (30, 31). Briefly, Fab, antibody, BSA, cell supernatant, and recombinant sCD93 functionalized beads were prepared by washing 10 μl of 3 µm diameter P(S/DVB) polystyrene bead 2× in 1 ml of PBS using a 6,000×g/1 min centrifugation. Beads were then suspended in 100 μl of cell-conditioned medium or 100 μl PBS + 10 μl of one of 12CA5 Fab, 50 mg/ml human IgG, 5% BSA, or recombinant sCD93, and incubated at room temperature with rotation for 30 min. Beads were then washed as above and suspended in 100 μl of PBS or serum-free DMEM. Apoptotic cell mimics were produced by preparing 4 μmol of a 20% phosphatidylserine, 79.9% phosphatidylcholine, 0.1% biotin-phosphatidylethanolamine mixture in chloroform. Subsequently, 10 μl of 3 µm diameter silica beads were added, the mixture vortexed for 1 min, and the chloroform evaporated under a stream of nitrogen for 1 hr. Beads were resuspended in 1 mL of PBS, vortexed, and washed as above. sCD93 covalently linked beads were prepared by washing 10 µL promag carboxy magnetic microspheres in 1 mL of MES at pH 7.2, and then suspended in 100 µL of the same buffer. To this, 1 mg of N-Ethyl-N’-(3-dimethylaminopropyl)carbodiimide was added and the beads incubated for 15 min on a rotator. The beads were washed twice in PBS and resuspended in 500 µL PBS to which 10 μL of 1% BSA or 10 μL of recombinant sCD93 were added and incubated for 4 hrs on a rotator at room temperature. Beads were washed and resuspended in 1 mL quenching solution (PBS with 40 mM glycine and 1% BSA), followed by a wash and resuspension in 100 μL storage buffer (0.1% BSA with PBS). All beads were stored at 4°C until used.

### Efferocytosis, Phagocytosis and Endocytosis Assays

These assays were performed as per our published methods (30, 31, 33). Briefly, for bead-based targets, 10 μL of the bead suspensions (∼10 beads:macrophage) were added to each well of macrophages in a 12-well plate. Apoptotic cells or bacterial targets were first opsonized by incubating 100 μl of the cell suspension with 10 μL of recombinant sCD93 or an equimolar concentration of BSA for 30 min at 37°C, followed by addition of the full 110 μL volume to a well of macrophages in a 12-well plate (∼10 apoptotic cells:macrophage, ∼50 microorganisms:macrophage). Once the targets were added, the samples were centrifuged at 200×g for 1 min to force contact between the macrophages and the targets, and the plate incubated at 37°C/5% CO_2_ for 30 min unless otherwise indicated. At the end of the experiment the media was aspirated, the samples washed vigorously 3 times with PBS to remove any unbound targets, and the cells fixed in 4% PFA for 20 min. If targets were biotinylated, a 1:500 dilution of fluorescent streptavidin was then added to detect uninternalized targets; for apoptotic Jurkat cells a 1:1000 dilution of FITC-conjugated anti-human CD3 was added to detect uninternalized targets. Cells were then washed 3 times with PBS, mounted on slides with Permafluor, and imaged as described above. Efferocytosis/phagocytosis was then quantified – for streptavidin-labeled samples the efferocytic index was quantified as the number of engulfed targets per macrophage, or as beads/cell if streptavidin staining was not available. For endocytosis assays, HA-CD93 expressing CHO cells were cooled to 10°C and labeled with a 1:100 dilution of 12CA5 Fab for 20 min. The cells were washed three times with PBS and then labeled for 20 min at 10°C with a 1:500 dilution of goat-anti-mouse Cy3-labeled Fab (non-crosslinking) or goat-anti-mouse Cy3-488 labeled F(ab’)_2_ (cross-linking). The cells were then washed with warmed serum-free RPMI, and placed in a 37°C incubator for 60 min. The cells were then fixed with PFA, any non-internalized CD93 labeled for 20 min with a 1:500 dilution of donkey-anti-goat Alexa488-labeled Fab, the samples washed 3 times in PBS, mounted on slides with Permafluor, and imaged as described above. Endocytosis was quantified using FIJI as the fraction of the Cy3-CD93 staining that lacked Alexa-488 colocalization.

### Suspension Efferocytosis Assay

Wild-type and shRNA expressing THP-1 monocytes were mixed in a 1:1 ratio, and 1 × 10^5^ of the mixed cells placed in a microfuge tube along with 1 × 10^6^ sCD93-coated beads. The mixture centrifuged at 300 × g/2 min to force contact between the cell and beads, and the tubes incubated for 20 min at 37°C. The supernatants were then removed and the cells fixed for 20 min with 4% PFA. The fixed cells were then placed into an imaging chamber and the number of beads bound by untransduced (zsGreen^−^) versus shRNA-transduced (zsGreen^+^) cells quantified, with at least 100 cells quantified per condition in each experiment.

### Recovery and Identification of the sCD93 Receptor

5×10^6^ THP-1 monocytes were deposited into each well of a 6-well tissue culture plate and differentiated into macrophages as described above. The cells were cooled to 10°C and to 3 wells 100 μl/well of BSA-conjugated promag carboxy magnetic microspheres were added, with the other 3 wells receiving 100 μl/well sCD93-conjugated microspheres. The samples were then centrifuged for 1 min at 200×g to force contact between cells and spheres, and the plate incubated for 10 min at 10 °C. Beads were then reversibly crosslinked to putative sCD93 receptors using the ReCLIP method (31, 34); briefly, the samples were washed with PBS and then PBS with 0.5 mM dithiobis[succinimidyl propionate] and 0.5 mM dithio-bismaleimidoethane added to the cells for 1 min at 10°C. The solution was then aspirated and the reaction quenched for 10 min with 5 mM L-cysteine and 20 mM Tris-Cl, pH = 7.4. The cells were then disrupted with RIPA buffer and the beads recovered with a magnetic column. The beads were then washed 3 times with RIPA buffer, vortexing the beads for 1 min between washes, and recovering the beads with a magnetic column. After the final wash the beads were suspended in 40 μl of Laemmli buffer with 50 mM DTT, boiled for 10 min, resolved on a 4%-20% SDS-PAGE gel for 4 hrs at 100V, and stained using a colloidal coomassie stain. Bands unique to the sCD93 beads were then excised using an Ettan Robotic Spot-Picker. Gel slices were destained with 50 mM ammonium bicarbonate and 50% acetonitrile, treated with 10 mM DTT and 55 mM iodoacetamide, and digested with trypsin. Peptides were extracted by a solution of 1% formic acid and 2% acetonitrile and freeze dried. Dried peptide samples were then suspended in a solution of 10% acetonitrile and 0.1% trifluoroacetic acid. Mass spectrometry analysis of each sample was performed using an AB Sciex 5800 TOF/TOF System. Raw mass spectrometry data analysis was performed using MASCOT database search (http://www.matrixscience.com).

### Immunoblotting

1 × 10^5^ shRNA-expressing THP-1 cells were removed from culture, washed once with PBS, and lysed in Laemmli’s buffer containing 10% β-mercaptoethanol and Halt protease inhibitor cocktail, boiled briefly, and then loaded onto a 10% SDS-PAGE gel. Gels were run for 30 min at 50 V, and then the voltage increased to 100 V until maximal migration. The separated proteins were then transferred to nitrocellulose membrane overnight at 40V and blocked for two hours with TBS-T containing 2.5% milk powder. Primary antibodies were added at a 1:250 dilution (CD44, β_1_, β_2_), or at 1:1000 (α_x_, GAPDH) for 2 hrs in TBS-T containing 2.5% milk powder, washed 3 × 15 min in TBS-T, and then an appropriate Dylight-700 labeled secondary antibody added at a 1:10,000 for 1 hr in TBS-T containing 2.5% milk powder. After 3 × 15 min washes in TBS-T the blots were imaged on an Odyssey CLx imaging system.

### Statistics and Data Analysis

All datasets were tested for normal distribution using a Shapiro-Wilk test. Normally distributed data is presented as mean ± SEM and analyzed using a Students T-test or ANOVA with Tukey correction. Non-parametric data is presented as median ± interquartile range and analyzed using a Kruskal-Wallis test with Dunn correction. All statistical analyses were performed using Graphpad Prism. All image analysis was performed using FIJI (29).

## Results

### CD93 Does Not Act as an Efferocytic or Tethering Receptor

Previous studies have suggested that CD93 acts as an efferocytic or tethering receptor, engaging targets either via the opsonin C1q, or directly (7, 10–14, 16, 35, 36). These claims are controversial, with studies indicating that C1q is not a CD93 ligand, and that the soluble form – not the membrane-bound form – of CD93 enhances efferocytosis (15, 16). As such, using multiple approaches we sought to directly assess whether the membrane-bound form of CD93 acts as an efferocytic or tethering receptor. First, CD93 was ectopically expressed in CHO cells, which have the molecular machinery required for phagocytosis and efferocytosis, but lack expression of phagocytic or efferocytic receptors (37). Transfection of CHO cells with the phagocytic receptor FcγRI rendered CHO cells highly phagocytic to beads opsonized with IgG, while transfection with the efferocytic receptor TIM4 rendered CHO cells efferocytic against apoptotic cells, serum-opsonized apoptotic cells, and apoptotic cell mimics (Figure 1A). In marked contrast, transfection with CD93 did not impart phagocytic or efferocytic activity against any of these targets (Figure 1A), indicating that CD93 is unlikely to function as an efferocytic or tethering receptor of either unopsonized or serum-opsonized apoptotic cells. We further explored the possibility that CD93 mediates efferocytosis via an opsonin by mimicking opsonin-mediated receptor cross-linking by oligomerizing HA-CD93 with anti-HA functionalized beads. No internalization of these bead was observed above the level of control beads coated with an irrelevant IgG (Figure 1B). Because CHO cells are not natively phagocytic or efferocytic, and may therefore be lacking a co-receptor or signaling molecule required by CD93, we repeated this experiment using macrophages which have previously been reported to be capable of CD93-dependent efferocytosis (16). To avoid Fc receptor activation, beads were functionalized with anti-HA Fab fragments or Fab fragments generated from an irrelevant IgG, rather than full-length antibodies. Anti-HA functionalized beads produced an accumulation of CD93 around the beads reminiscent of a phagocytic cup (Figure 1C), but did not induce bead engulfment (Figure 1D). Next, we quantified CD93 endocytosis following antibody-mediated cross-linking of CD93, as previous reports have demonstrated that cross-linking is sufficient to activate and induce the endocytosis of phagocytic and efferocytic receptors via the same signaling pathways triggered by *bona fide* phagocytic/efferocytic targets (33, 38–40). Cell-surface HA-CD93 was labeled under cross-linking (Cy3-F(ab’)_2_) or non-crosslinking (Cy3-Fab) conditions, endocytosis allowed to occur for 30 min, and residual surface Fab/F(ab’)_2_ labeled with Alexa-488. Under these conditions, endocytosed CD93 is labeled only with Cy3, while non-endocyotsed CD93 is labeled with both Cy3 and Alexa-488. Although cross-linking resulted in significant clustering of CD93 on the cell surface (Figure 1E), CD93 was endocytosed at an equal rate in non-crosslinked and cross-linked samples (Figure 1F). Finally, no tethering of apoptotic Jurkat cells was observed by non-efferocytic HeLa cells ectopically expressing CD93 (Figure 1G). Combined, these data demonstrate that the transmembrane form of CD93 is not an efferocytic or tethering receptor.

**Figure 1:**
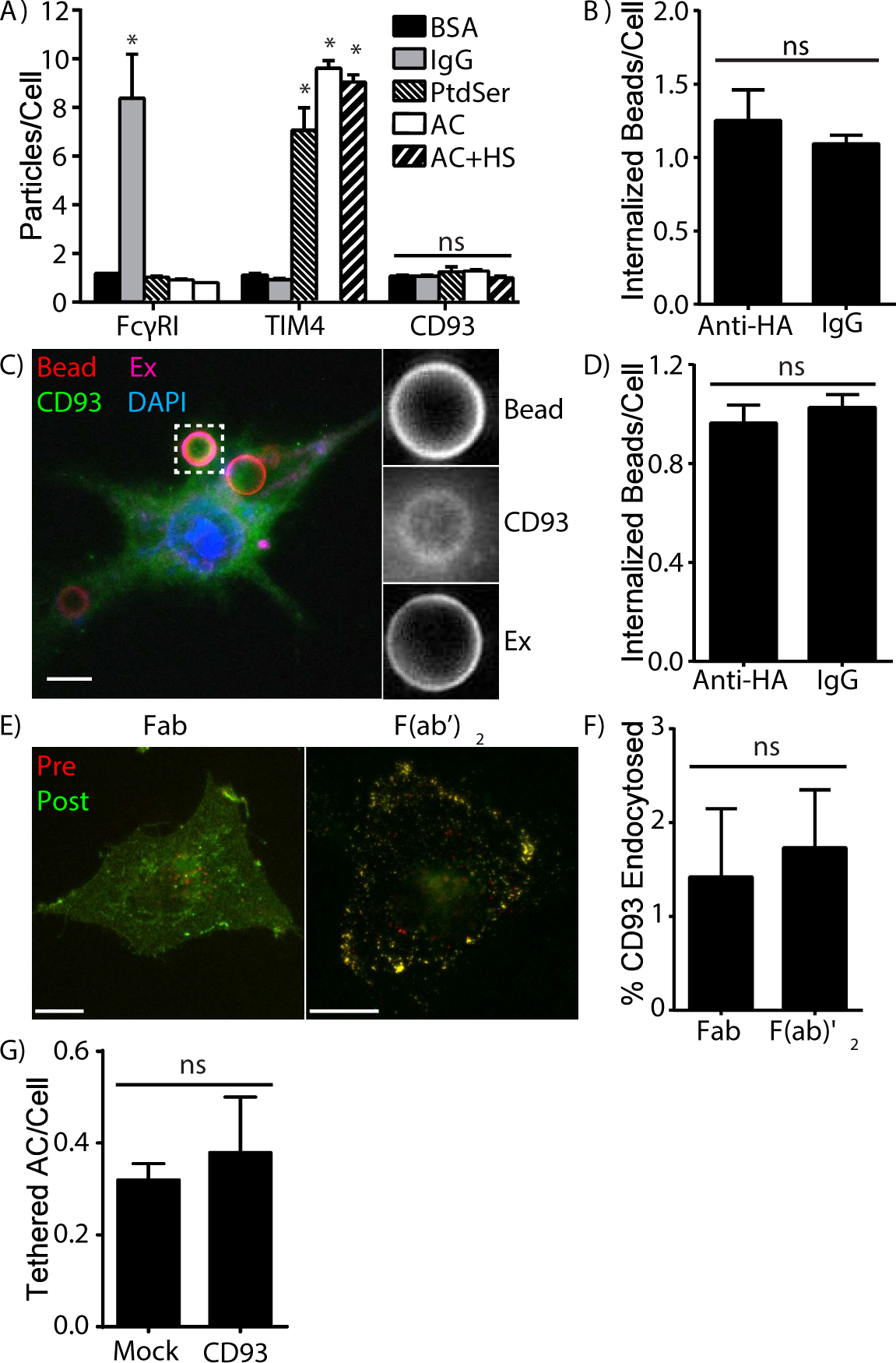
CD93 is not a phagocytic or efferocytic receptor. **A)** Total bound and engulfed control beads (BSA), phagocytic targets (IgG coated beads/IgG), apoptotic cell mimics (PtdSer), apoptotic cells (AC) and human-serum opsonized apoptotic cells (AC+HS) by CHO cells ectopically expressing human CD93, FcγRI and TIM4. **B)** Phagocytosis of beads functionalized with anti-HA or an irrelevant IgG by CHO cells ectopically expressing HA-tagged CD93. **C)** Accumulation of CD93 beneath anti-HA Fab functionalized beads in RAW 264.7 macrophages ectopically expressing HA-CD93-GFP. **D)** Uptake of beads functionalized with anti-HA Fabs or an irrelevant IgG by RAW 264.7 cells ectopically expressing HA-tagged CD93. **E-F)** Images (E) and quantification (F) of non-crosslinked (Fab) and crosslinked (F(ab’)2) HA-CD93 ectopically expressed in CHO cells. Cell-surface CD93 is labeled before (Pre, red) and after (Post, green) endocytosis. **G)** Number of tethered apoptotic Jurkat cells by mock-transfected and CD93-transfected HeLa cells. Data is representative of, or quantifies as mean ± SEM, a minimum of 50 images acquired over a minimum of 3 independent experiments. * = p < 0.05, ns = p > 0.05, ANOVA with Tukey correction (A, compared to BSA for the same receptor), or Students *t*-test (B,D,F). Scale bars are 5 µm.

### CD93 Is Shed During Inflammation and Macrophage Differentiation

Soluble CD93 has been identified in human serum and is produced following stimulation of human and murine myeloid cells with inflammatory stimuli (4, 28). CD93 is expressed on circulating myeloid cells, but not on circulating lymphoid cells, and consistent with this we found that ∼60% of human PBMCs expressed CD93 (Figure 2A). This CD93 was shed upon stimulation with PMA, with permeabilization prior to immunostaining confirming that PMA stimulation induced shedding rather than endocytosis (Figure 2A). As PMA is a non-physiological stimulus, we assessed CD93 shedding by human monocytes following LPS and TNFα stimulation, with both ligands inducing a time-dependent shedding of CD93 (Figure 2B). In murine cells CD93 is shed by an uncharacterized matrix metalloproteinase (MMP) (4). Unexpectedly, the pan-MMP inhibitor GM6001 did not inhibit PMA-induced CD93 shedding by human PBMCs, whereas TAPI2 – which inhibits both MMP and a disintegrin and metalloproteinase (ADAM) family proteases – prevented PMA-induced CD93 cleavage (Figure 2C). Next, we assessed CD93 expression during differentiation of primary human monocytes into non-inflammatory (M0 and M2) and inflammatory (M1) macrophages. CD93 was downregulated early during macrophage differentiation (Figure 2D), and was absent from fully matured M0, M1, or M2 macrophages (Figure 2E). Lastly, we assessed the shedding of CD93 following apoptosis, as both ADAMs and MMPs are active during apoptosis (41, 42), with staurosporine-induced apoptosis of PBMC’s resulting in the TAPI2-inhibitable shedding of CD93 (Figure 2F). Combined, these data further argue against CD93 acting in a membrane-bound efferocytic or phagocytic receptor, as receptor shedding during infiltration of inflamed sites, and during macrophage differentiation, would deprive myeloid cells of an efferocytic receptor in environments where they would be expected to encounter apoptotic cells. As such, it is most probable that CD93 mediates efferocytosis in its soluble form.

**Figure 2:**
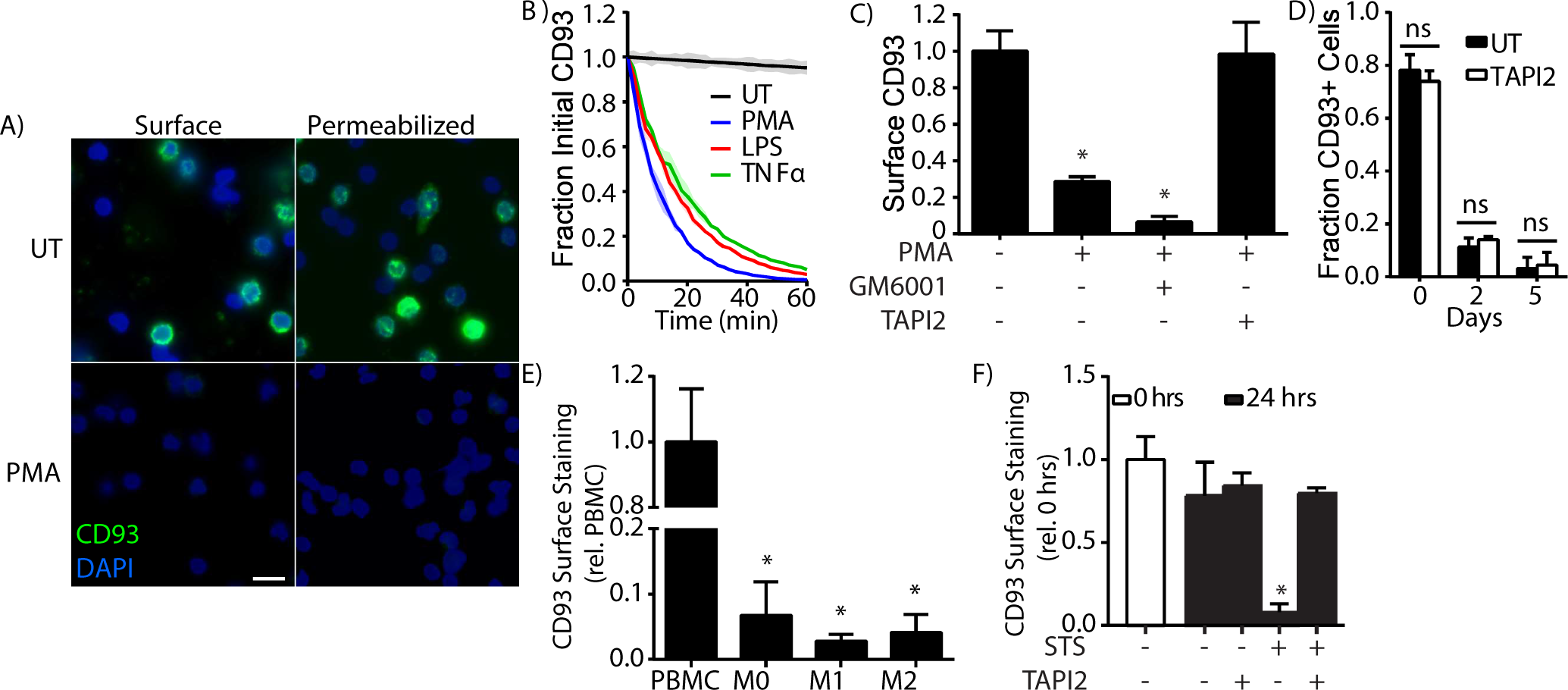
CD93 is shed during inflammation and macrophage differentiation. CD93 cleavage and expression was quantified in primary human monocytes and macrophages using immunofluorescence microscopy. **A)** Cell surface (Surface) and whole-cell (Permeabilized) CD93 immunostaining on PBMCs left untreated (UT) or stimulated with 100 ng/mL PMA for 1 hr. Cells are counter-stained with DAPI. **B)** Time-course of CD93 cleavage from monocytes following stimulation with 100 ng/mL PMA, 10 ng/ml LPS, or 100 ng/ml TNFα. **C)** Effect of the MMP inhibitor GM6001 and MMP/ADAM inhibitor TAPI2 on PMA-induced CD93 cleavage by human monocytes. **D)** CD93 surface expression on human PBMC’s at day 0, 2 and 5 of PBMC-to-M0 macrophage differentiation. **E)** Intensity of CD93 staining on M0, M1 and M2 macrophages, measured relative to undifferentiated PBMCs. **F)** CD93 shedding following 24 hrs treatment of PBMC’s with staurosporine (STS) ± TAPI-2, relative to CD93 on freshly isolated PBMCs (0 hrs). Data is representative of (A) or quantified as mean ± SEM (B–F), ≥17 cells imaged over 3 independent experiments. * = p < 0.05 compared to untreated (C), PBMC (E) and 0 hrs (F), ns = p > 0.05, ANOVA with Tukey correction. Scale bar is 10 μm.

### sCD93 Functions as an Opsonin

Based on previous studies which demonstrate that sCD93 enhances efferocytosis, and our data indicating that membrane-bound CD93 does not directly function as an efferocytic or tethering receptor (Figure 1), we next assessed whether sCD93 functions as an opsonin. First, we coated 3 µm diameter beads with supernatants from unstimulated versus PMA-stimulated monocytes, and then assessed the phagocytic activity of J774.2 macrophages against these targets. Mimics opsonized with supernatants from PMA-stimulated cells were engulfed at a higher rate than beads opsonized with media from unstimulated cells, with depletion of sCD93 abrogating this effect (Figure 3A). Next, apoptotic cell mimics (30) comprised of silica beads coated in a lipid mixture which mimics the plasma membrane of apoptotic cells (20% phosphatidylserine, 80% phosphatidylcholine) were coated with purified recombinant sCD93 or a control protein (BSA), with sCD93 modestly increasing bead binding by macrophages (Figure 3B). As sCD93 appeared to be functioning as an apoptotic cell opsonin, we next assessed the capability of recombinant sCD93 to opsonize apoptotic Jurkat cells. sCD93 significantly increased both the fraction of macrophages engulfing at least one apoptotic cell (Figure 3C), and the number of apoptotic cells engulfed per macrophage (Figure 3D), resulting in a net 4-fold increase in efferocytosis. No uptake of BSA- or sCD93-opsonized non-apoptotic cells was observed (data not shown). Finally, the opsonic effect of sCD93 was assessed against a range of microorganisms, using a panel of organisms representing a broad range of microbial diversity and capsulation states. No opsonic effect was observed with any of the tested Gram-negative bacteria (Figure 3E), Gram-positive bacteria (Figure 3F) or fungi (Figure 3G), demonstrating that sCD93 functions as an apoptotic cell-specific opsonin.

**Figure 3:**
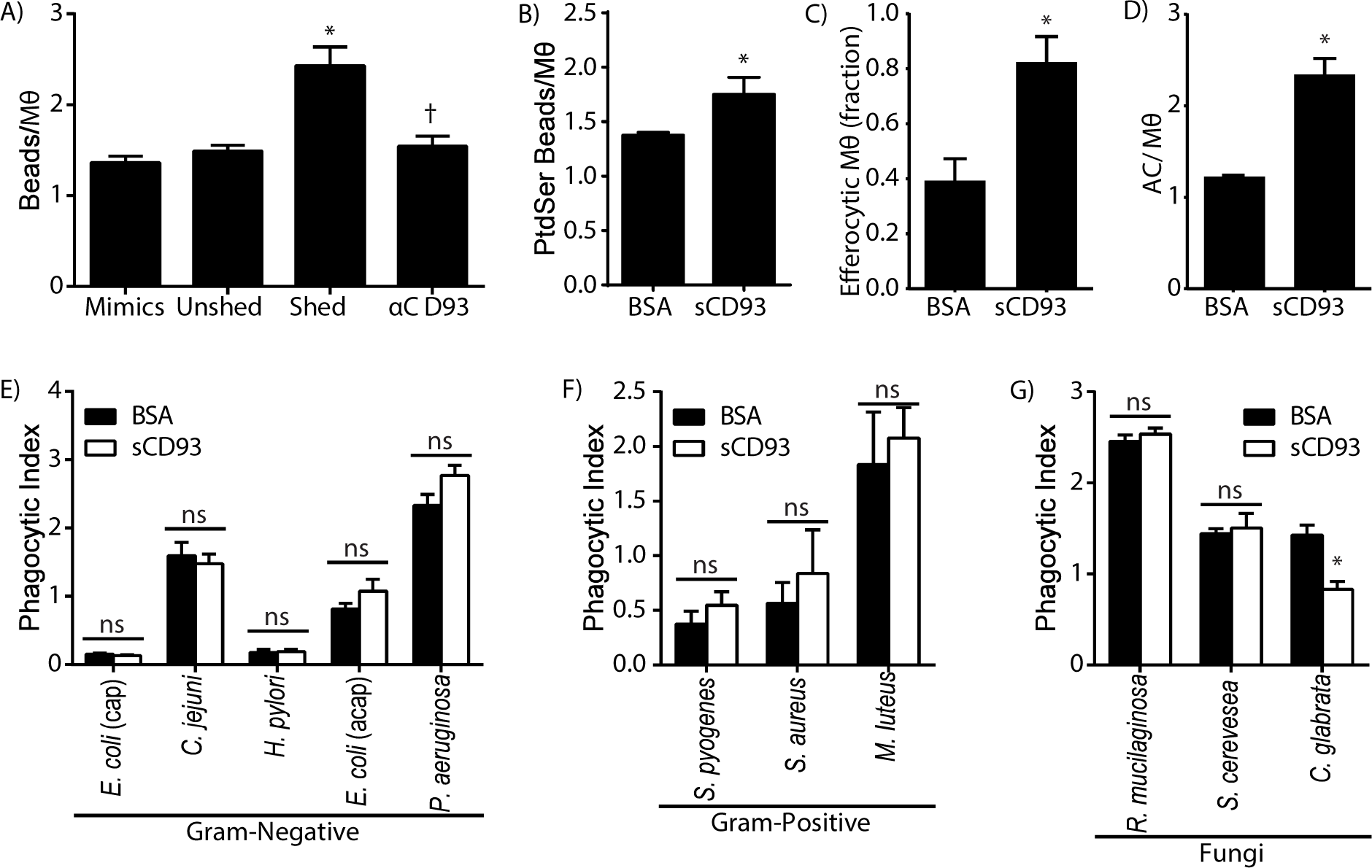
CD93 is an apoptotic cell opsonin. **A)** Number of supernatant-opsonized apoptotic cell mimics engulfed by J774.2 macrophages (Mθ), using mimics opsonized with cell culture media (Media), monocyte-conditioned medium (Unshed), supernatants from PMA-activated monocytes (Shed), or CD93-depleated supernatants from PMA-activated monocytes (αCD93). **B)** Total number of recombinant sCD93-coated versus BSA-coated apoptotic cell mimics bound or engulfed by J774.2 macrophages. **C)** Fraction of macrophages that engulfed one or more BSA-opsonized or sCD93- opsonized apoptotic cells. **D)** Average number of BSA-opsonized versus sCD93-opsonized apoptotic cells (AC) engulfed per J774 macrophage over 1 hr. **E-G)** Phagocytosis of unopsonized and sCD93 opsonized (E) Gram-negative bacteria: capsulated (cap) and acapsulated (acap) *Escherichia. coli*, capsulated *Campylobacter jejuni* and *Heliobacter pylori*, and acapsular *Pseudomonas aeruginosa*; (F) Gram-positive bacteria: capsular *Streptococcus pyogenes* and *Staphylococcus aureus*, and acapsular *Micrococcus luteus*; and (G) fungi: capsular *Rhodotorula mucilaginosa*, and acapsular *Sacchoarmyces cerevesea* and *Candida glabrata*. n = 3, mean ± SEM. * p < 0.05 compared to unopsonized, † p < 0.05 compared to sCD93 opsonized, ns = p > 0.05 compared to non-opsonized; ANOVA with Tukey correction (A), Students *t*-test (B-D), or paired *t*-test comparing the same organism ± sCD93 (E-G).

### Identification of the sCD93 Receptor

We next used a ligand-precipitation approach to identify the sCD93 receptor on macrophages. Briefly, sCD93 was covalently conjugated to magnetic beads, these beads centrifuged onto macrophages, a reversible cross-linking reagent used to stabilize the sCD93-receptor interaction, and the CD93-receptor complex recovered with a magnetic column (31, 34). sCD93-coated beads precipitated two unique bands not observed in the control, both of which were much larger than sCD93 (68 kDa, Figure 4A). These bands were excised from the gel and the constituent proteins identified by liquid-chromatography/mass spectrometry. While many proteins were non-specifically cross-linked to the sCD93 beads, filtering of the dataset for high abundance proteins found in at least 3 of 4 independent experiments identified only 8 proteins as putative sCD93 receptor components (Table 1). These included three phagocytic integrin subunits (α_x_, β_2_ and β_1_), CD44 which regulates integrin-mediated phagocytosis, and several integrin-associated signaling molecules, indicating that the sCD93 receptor is likely an integrin (43–46). To determine which of these proteins function as the sCD93 receptor, α_x_, β_1_, β_2_, and CD44 deficient THP-1 cell lines were generated using lentiviral-shRNAs (Figure 4B). Efferocytosis assays were then performed using sCD93 coated beads and THP-1 cells in suspension, as differentiation of THP-1 cells to adherent macrophages selected for cells with partial integrin knockdown (data not shown). shRNAs against α_x_ and β_2_ decreased both the fraction of macrophages engaging in efferocytosis and the number of beads efferocytosed per cell (Figure 4C-D), with either shRNA producing a net decrease in efferocytosis of ∼88%. Knockdown of β_1_ or CD44 did not impair sCD93-mediated efferocytosis, thus identifying α_x_β_2_ integrin as the receptor for sCD93.

**Figure 4:**
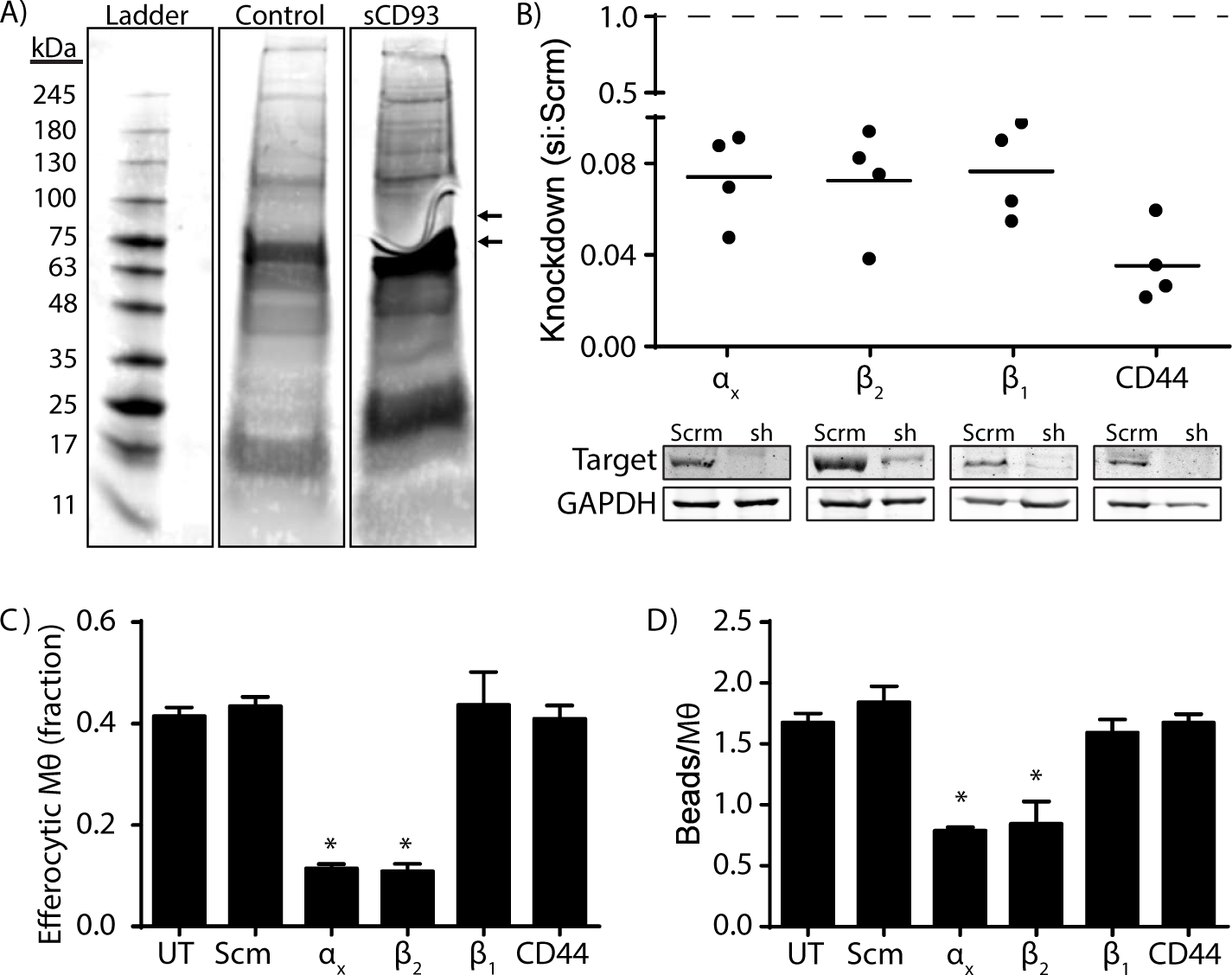
Identification of the sCD93 receptor. **A)** Representative blot demonstrating the precipitation of putative sCD93 receptors with sCD93-functionalized versus BSA-functionalized (Control) magnetic beads and reversible cross-linking. Arrows indicate protein bands uniquely precipitated by the sCD93- functionalized beads. **B)** Confirmation of shRNA knockdown of putative CD93 receptors in THP-1 macrophages. Knockdown levels are expressed as fold-knockdown after normalization to GAPDH staining. Representative immunoblots are shown below the graph. Scrm = THP-1 expressing scrambled (control) shRNA, sh = shRNA-expressing THP-1 cells. Dashed line indicates the expression level in Scrm-siRNA expressing cells. **C)** Effect of shRNA knockdown of putative sCD93 receptors on the fraction of macrophages engulfing one or more sCD93-opsonized beads. **D)** Effect of shRNA knockdown on the number of sCD93-opsonized beads engulfed per macrophage. Data is representative of (A) or quantifies (B-D) four independent experiments. Data is presented as mean ± SEM. * p < 0.05, ANOVA with Tukey correction.

**Table 1:** MS/MS Identification of Putative Members of the sCD93 Receptor Complex

| **Protein** | **Symbol** | **Peptides\*** | **Coverage (%)\*\*** |
| --- | --- | --- | --- |
| *Integrins* |  |  |  |
| Integrin $\alpha_x$ | ITGAX (CD11c) | 6 | 10.8 |
| Integrin $\beta_1$ | ITGB1 (CD29) | 12 | 24.3 |
| Integrin $\beta_2$ | ITGB2 (CD18) | 19 | 31.1 |
| *Integrin-Associated Phagocytic Receptors* |  |  |  |
| CD44 | CD44 | 3 | 4.6 |
| *Integrin-Associated Signaling Molecules* |  |  |  |
| Talin | TLN1 | 48 | 29.3 |
| Focal Adhesion Kinase 2 | FAK2 | 27 | 33.2 |
| PAK2 | PAK2 | 8 | 20.4 |
| Moesin | MOES | 49 | 88.2 |

\* Peptides = total number of unique peptides from each protein identified across four independent experiments.
\*\* % Coverage = portion of protein sequence covered by the identified peptides.

### Identification of the Ligand and Receptor Binding Domains in sCD93

The identification of an integrin as the sCD93 receptor was unexpected, as human CD93 lacks a canonical RGD or other integrin-recognition motif (47). As such, we identified the sCD93 receptor-binding and apoptotic cell-binding domains of CD93 using deletion mutagenesis (Figure 5A), preparing purified FLAG-tagged sCD93 mutants lacking the CTLD (CD93^ΔCTLD^), the EGF-like repeats (CD93^ΔEGF^), or the mucin domain (CD93^ΔMucin^). First, efferocytosis assays were performed using beads non-specifically coated in these mutant proteins, allowing us to probe sCD93 binding by macrophages independently of sCD93’s ligand-binding activity. These experiments demonstrated that the EGF-like domains are required for sCD93 binding by THP-1-derived macrophages, indicating this domain is recognized by α_x_β_2_ (Figure 5B). Next, efferocytosis of apoptotic cells opsonized with these mutant proteins was assessed; this confirmed the requirement for the EGF-like domains, and in addition, demonstrated that the opsonic effect of sCD93 was lost when the CTLD domain was deleted (Figure 5C). Combined, these results indicate that CD93 acts as a bipartite opsonin; binding to apoptotic cells via its CTLD, and then bridging the apoptotic cell to α_x_β_2_ integrin on phagocytes via its EGF-like domains.

**Figure 5:**
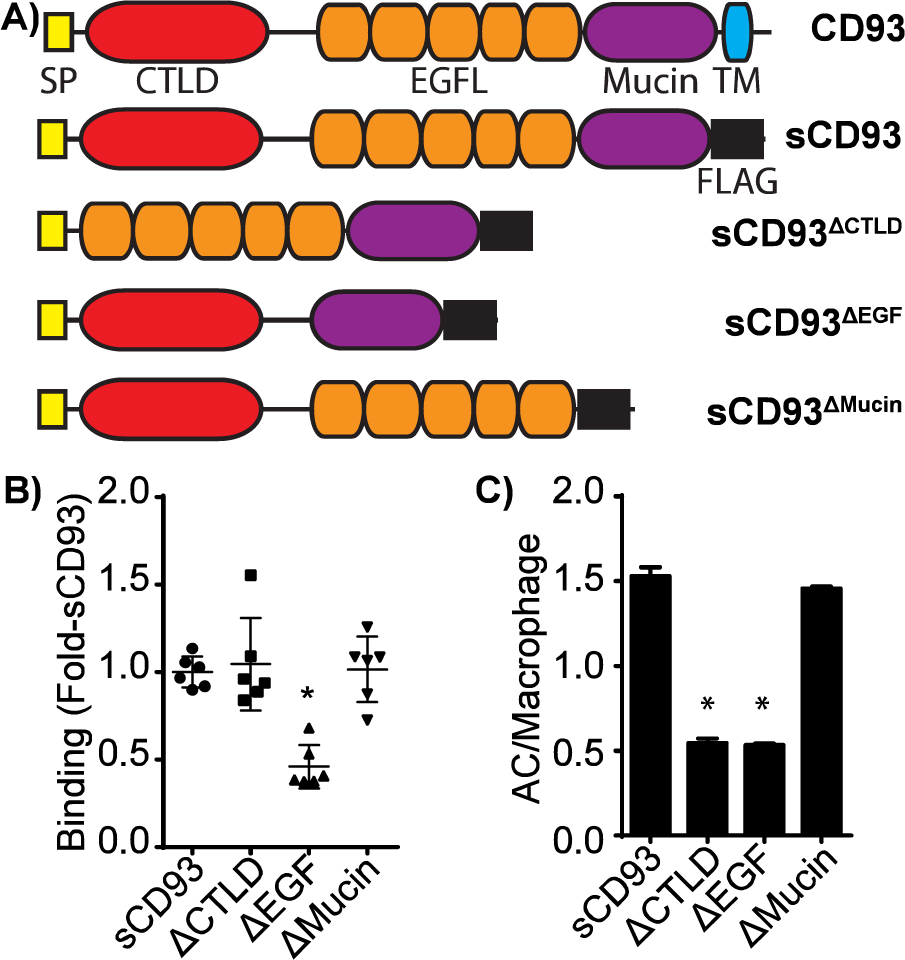
Identification of sCD93-receptor and apoptotic cell binding domains in sCD93. **A)** Structural diagrams of FLAG-tagged full-length and CTLD-deficient (sCD93^ΔCTLD^), EGFL-deficient (sCD93^ΔEGF^) and Mucin-deficient (sCD93^ΔMucin^) mutants of sCD93. SP = signal peptide, CTLD = C-Type Lectin-Like Domain, EGFL = EGF-like, TM = transmembrane domain. **B)** Binding of beads opsonized with full-length and mutant sCD93 by THP-1 macrophages. **C)** Binding of apoptotic cells (AC) by full-length and mutant sCD93. Data quantifies three independent experiments. Data is presented as mean ± SEM, * p < 0.05, Kruskal-Wallis with Dun Correction (B) or ANOVA with Tukey Correction (C).

## Discussion

In this study, we have demonstrated that soluble CD93 is rapidly released from monocytes by proteolytic cleavage in response to inflammatory stimuli and during monocyte-to-macrophage differentiation. The soluble fragment then acts as an apoptotic cell opsonin, binding to apoptotic cells via its CTLD domain and bridging the apoptotic cell to α_x_β_2_ integrins on phagocytes via its EGF-like domain. This opsonic activity greatly enhances the uptake of apoptotic cells by phagocytes, to an extent similar to that observed for other apoptotic cell opsonins (48–52), and could be abrogated by knockdown of the α_x_β_2_ integrin, or by deletion of the apoptotic cell-binding or integrin-binding domains of CD93. Combined, these data are consistent with a model in which CD93 is shed during inflammation – both following the extravasation of leukocytes into the inflamed sites, and during the differentiation of monocytes into macrophages – thus providing the localized delivery of an apoptotic cell opsonin to environments typified by high levels of cell death.

CD93 is a member of the group XIV C-type lectin family, which also includes thrombomodulin, CLEC14A and CD248 (53). While all members of this family contain a C-type lectin-like domain, carbohydrate binding has not been reported for any of these receptors. Rather, all known ligands for this gene family are proteins – thrombomodulin binds thrombin (54), while the extracellular matrix protein multimerin-2 was recently shown to be a ligand for CLEC14A, CD248 and CD93 (55). The CD93- multimerin-2 interaction has been characterized in depth, with CD93 binding a coiled-coiled domain in multimerin-2 via its CTLD (56). Consistent with this, we observed minimal opsonization of beads coated in a lipid mixture that mimics the plasma membrane of an apoptotic cell, while robust opsonization of apoptotic cells was observed. This would suggest that CD93 is not recognizing the common “eat-me” signal phosphatidylserine, and instead is recognizing another – potentially proteinaceous – “eat me” signal. Multimerin-2 is unlikely to be this ligand, as multimerin-2 is not expressed by the target cell type used in this study, nor is it upregulated during apoptosis (57–59). Previous studies have suggested that C1q and related lectins (MBL, SPA) may also be CD93 ligands (7, 13, 14, 35, 36), and C1q is known to opsonize apoptotic cells (60), suggesting that C1q may represent a putative ligand for CD93 on apoptotic cells. This, however, is unlikely as the reported interaction between CD93 and C1q is thought to be artefactual with CD93-deficient macrophages retaining normal phagocytic activity to C1q-opsonized targets, and attempts to directly measure C1q-CD93 interactions failing to identify an interaction between these proteins (15, 16). Consistent with these studies, we observed sCD93-dependant opsonization in the absence of serum, further suggesting that C1q is not a ligand for sCD93. Several proteinaceous “eat me” signals have been reported in the literature that may act as sCD93 ligands. These include normally intracellular proteins which are trafficked to the cell surface or secreted upon induction of apoptosis (e.g. calreticulin, tubby and TULP1), serum-derived and macrophage-secreted pattern recognition receptors (e.g. PTX3, Gal3), and alterations in post-transcriptional glycosylation that may expose protein interaction motifs normally masked by carbohydrate groups (51, 52, 61–64). Finally, it is possible that the CTLD of CD93 retains some carbohydrate-binding capability, and thus CD93 may directly recognize the aberrant glycosylation patterns found on apoptotic cells (63).

α_x_β_2_ (also known as complement receptor 4, and as CD11c/CD18) has a well-established role as a phagocytic receptor that mediates the engulfment of iC3b-opsonized bacteria and apoptotic cells (65, 66). The signaling and function of this integrin is well understood, and indeed, our precipitation of the sCD93 receptor identified many of the proteins known to regulate α_x_β_2_-mediated phagocytosis as part of the sCD93 receptor complex. Talin induces integrin conformational changes from a low-affinity to high-affinity conformation (67), focal adhesion kinase mediates the outside-in signaling of integrins (68, 69), while PAK2 is activated by focal adhesion kinase to mediate cytoskeletal rearrangements downstream of integrin activation (70, 71). Moreover, the presence of β_1_ integrin and CD44 in our immunoprecipitates, despite their lack of activity as a sCD93 receptor, is likely a result of the structuring of integrins and their co-receptors into focal contacts; pre-formed assemblies of multiple integrins, their co-receptors, and signaling molecules (72, 73). This clustering enhances integrin function through increased avidity and expands the diversity of ligands that can be engaged by an activated cluster; however, this pre-clustering also renders all proteins in the cluster to become susceptible to the proximity-based crosslinking method used in this study (34, 74–76). The identification of an integrin as the sCD93 receptor was initially unexpected due to the absence of a canonical integrin binding motif in CD93. However, α_x_β_2_ is known to bind proteins lacking these motifs, often by binding EGF-like domains in the target protein, consistent with our identification of the EGF-like domains of sCD93 as the α_x_β_2_ binding site (77, 78).

Previous investigations into the role of CD93 in leukocyte function have identified several seemingly disparate roles for sCD93 which may be explained by our identification of α_x_β_2_ integrin as the sCD93 receptor. Jeon *et al.* demonstrated that sCD93 induced monocyte-to-macrophage differentiation and increased TNFα synthesis following LPS exposure (9). Engagement of β_2_ integrins is known to induce monocyte-to-macrophage differentiation through PCK-dependent suppression of the transcriptional repressor Foxp1 and concurrent upregulation of the M-CSF receptor (79, 80). While this phenomenon was observed downstream of α_m_β_2_, all β_2_-integrins activate the same PKC signaling pathway via their common β_2_ chain – and thus α_x_β_2_ would be expected to activate the same macrophage differentiation program as α_m_β_2_ (81–83). Similarly, β_2_-integrins enhance TNFα release following LPS stimulation. Endocytosis of integrins – a likely outcome following engaging a soluble ligand such as sCD93 – enhances LPS responses by co-endocytosing TLR4, thus increasing TLR4 signaling from endosomes (84–86). Moreover, TNFα release is further enhanced through the cooperative signaling of the integrin signaling molecule FAK and the TLR signaling molecule MyD88, an interaction which has been previously shown to synergistically increase TNFα release following LPS stimulation (87). In another report, CD93-deficiency was associated with enhanced leukocyte recruitment in a model of peritonitis, with recruitment reduced to normal levels in CD93 chimeras (27). α_x_β_2_ is known to be a potent regulator of monocyte adhesion to endothelial-expressed ICAM-1, ICAM-4 and VCAM-1, and is required for monocyte recruitment to the inflamed peritoneum (88, 89). It is possible that CD93 may act as a competitive inhibitor of α_x_β_2_, thus decreasing intravascular adhesion and limiting leukocyte infiltration into inflamed tissues.

Developing B cells represent another immune cell population which express CD93. CD93 expression, as detected by the AA4.1 and 493 antibodies, is a classical marker of developing B cells (90) and demarcates hematopoietic stem cells, B cell/myeloid bipotential progenitors, and all stages of B cell development up to the transitional B cells which migrate from the bone marrow to become naive B cells in peripheral lymphoid tissues (90, 91). Although the role of CD93 in B cell development is unclear, it is established that CD93-deficiency does not prevent B cell development (16), but interestingly, diabetes-susceptible NOD mice bear a mutation in CD93 that limits CD93 expression during B cell lymphopoiesis (18, 92). B cell development is characterized by high levels of apoptosis, with ∼90% of lymphoid progenitors that commit to the B cell lineage undergoing apoptosis prior to developing into naïve B cells (93). While the receptors involved in removing apoptotic cells from the bone marrow have not been well characterized, our data suggest that CD93 released from apoptosing B cells may enhance efferocytosis in the bone marrow in an autocrine or paracrine manner. Poor clearance of these cells may promote autoimmunity, as failed efferocytosis of apoptotic B cells has been suggested to increase the stimulation of immature self-reactive B cells, thereby allowing them to undergo sustained receptor editing or to bypass central tolerance (94). Moreover, CD93 is re-expressed during plasma cell differentiation, which is another phase of the B cell lifecycle typified by high levels of apoptosis (95), further consistent with a model in which B cells release sCD93 to ensure their timely removal during developmental stages where B cell apoptosis is common. Consistent with this hypothesis, MMP and ADAM proteases are known to be activated during apoptosis (41, 42), and we observed shedding of CD93 in response to staurosporine-induced apoptosis of PBMCs.

In addition to its multifaceted effects on myeloid cells and opsonic activity, CD93 also has an established role as a potent angiogenic factor (96). The mechanism by which CD93 promotes angiogenesis has not been fully elucidated, with evidence supporting the presence of two distinct pathways. The first putative pathway is regulated by sCD93, with sCD93 directly inducing endothelial cell division, migration and tubule formation with a potency similar to that of VEGF (96). The endothelial receptor which recognizes sCD93 is unknown, but is unlikely to be α_x_β_2_ as this integrin is not normally expressed by endothelium (97, 98). The second angiogenic pathway enhances endothelial cell migration, is driven by full-length (membrane-bound) CD93 on endothelium, and requires interactions between membrane-bound CD93 and dystroglycan or multimerin-2 in the extracellular matrix (24, 55, 56). The signaling downstream of the CD93-multimerin-2 interaction has not been characterized, however, CD93-dystroglycan interactions induce phosphorylation of the intracellular tail of CD93, with this phosphorylation required for actin reorganization and endothelial cell migration (24). How CD93 induces cell migration remains unknown, but CD93 is known to interact with moesin and GIPC, suggesting these proteins may play a role in mediating signaling and cell migration through full-length CD93 (22, 23). The dual-role of sCD93 as an angiogenic factor and apoptotic cell opsonin suggests that CD93 plays an important role in the resolution of inflammation, as the removal of apoptotic cells and revascularization of tissues are key steps during the resolution of inflammation, and are required to return the tissue to a healthy state.

We conclude that CD93 is shed from myeloid cells under inflammatory conditions, with the resulting sCD93 opsonizing apoptotic cells and enabling efferocytosis via α_x_β_2_, while simultaneously enhancing angiogenesis within the inflamed site. This mechanism would ensure the concentrated, localized delivery of sCD93 to the inflamed environment – thus enabling the ready removal of cells killed during inflammatory responses, and preparing the inflamed site for revascularization following resolution of inflammation. More generally, CD93 may be shed by multiple cell types in response to cell death, thus allowing these cells to ensure their removal through autocrine or paracrine opsonization.

## Acknowledgements

The authors would like to thank Dr. Carole Creuzenet and Jocelynn Peters (University of Western Ontario) for culturing and preparing the *Campylobacter jejuni* and *Helicobacter pylori* used in this study. This study was funded by Canadian Institutes of Health Research Operating Grant MOP-123419, Natural Sciences and Engineering Research Counsel of Canada Discovery Grant 418194, and an Ontario Ministry of Research and Innovation Early Research Award to BH. JDD was funded by a Canadian Institutes of Health Research Operating Grant MOP-286719 whereas ENP was funded by an Alexander Graham Bell Doctoral Canada Graduate Scholarship from the Natural Sciences and Engineering Council. The funding agencies had no role in study design, data collection and analysis, decision to publish, or preparation of the manuscript.

---

**Supplemental Table 1:**
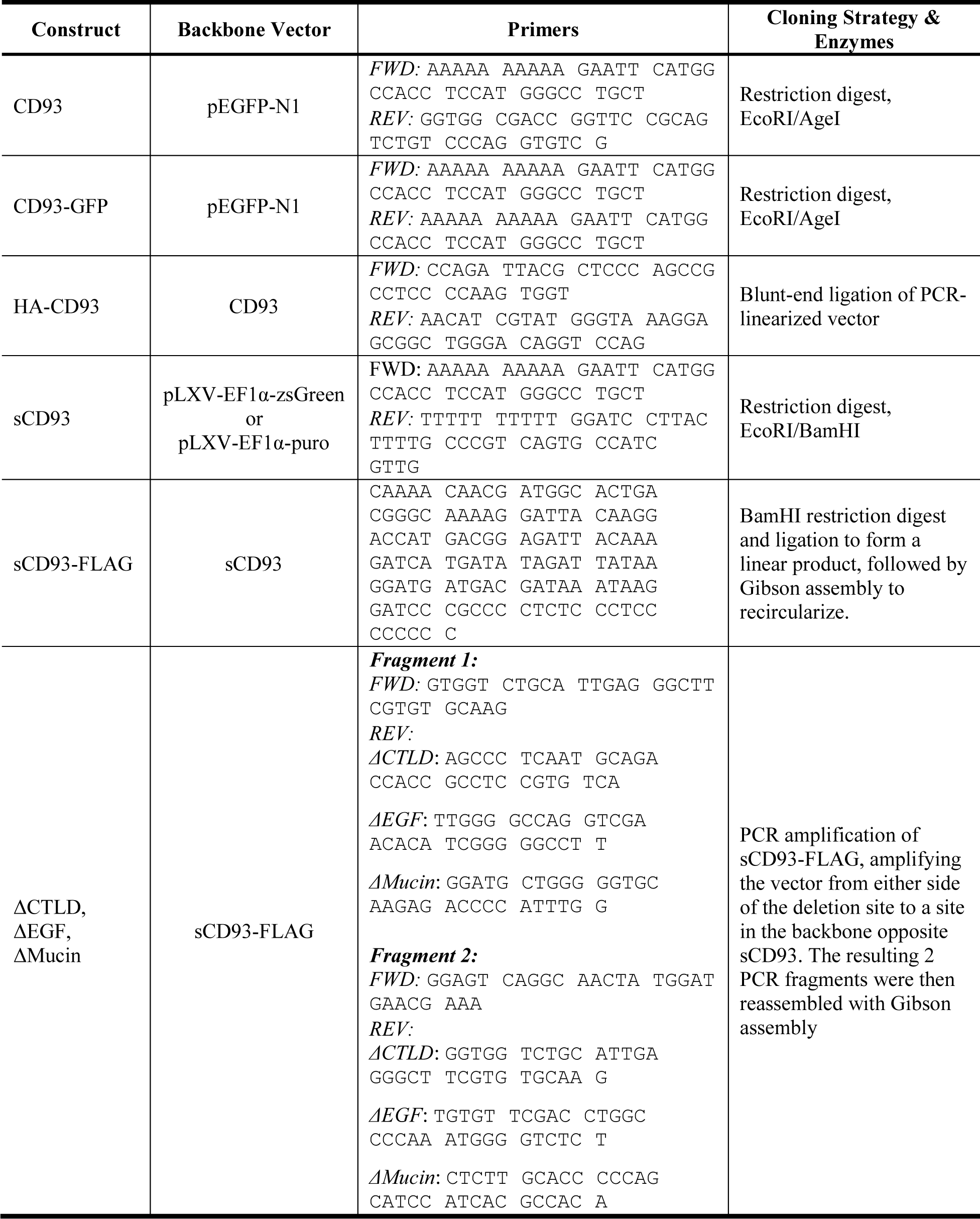
Enzymes and cloning approaches used to prepare all expression vectors used in this study. *FWD* = forward primer, *REV* = reverse primer.

**Supplemental Table 2:**
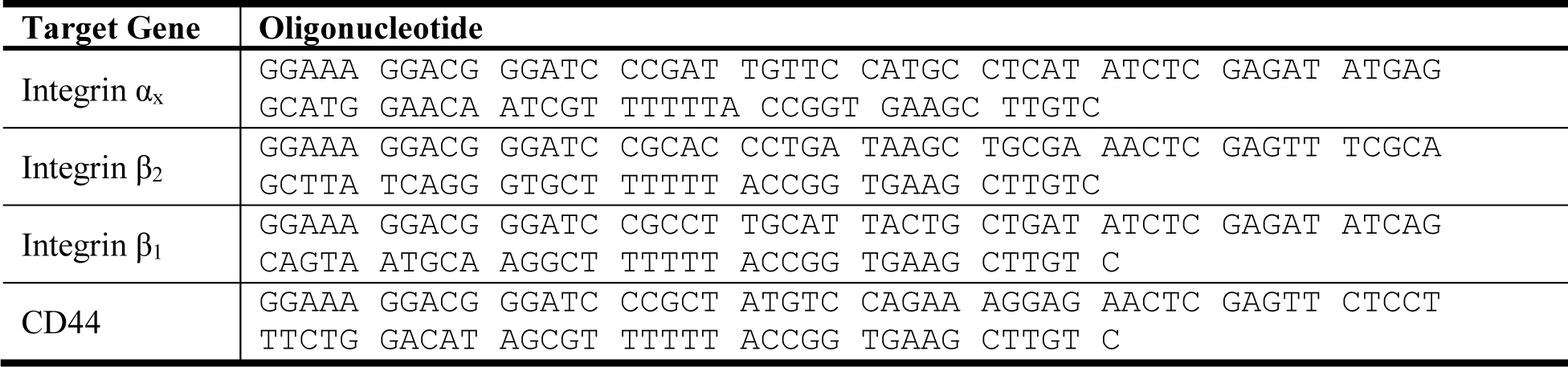
shRNA vector sequences.

